# Exome-capture RNA-sequencing of decade-old breast cancers and matched decalcified bone metastases identifies clinically actionable targets

**DOI:** 10.1101/120709

**Authors:** Nolan Priedigkeit, Rebecca J. Watters, Peter C. Lucas, Ahmed Basudan, Rohit Bhargava, William Horne, Jay K. Kolls, Zhou Fang, Margaret Q. Rosenzweig, Adam M. Brufsky, Kurt R. Weiss, Steffi Oesterreich, Adrian V. Lee

**Affiliations:** Department of Pharmacology and Chemical Biology, University of Pittsburgh, Pittsburgh, Pennsylvania; Women’s Cancer Research Center, University of Pittsburgh Cancer Institute, Pittsburgh, Pennsylvania; Department of Orthopedic Surgery, University of Pittsburgh, Pittsburgh, PA; Magee-Women’s Research Institute, Magee-Women’s Research Hospital of University of Pittsburgh Medical Center, Pittsburgh, Pennsylvania; Department of Human Genetics, University of Pittsburgh, Pittsburgh, PA; Department of Pathology, University of Pittsburgh Medical Center, Pittsburgh, Pennsylvania; Richard King Mellon Foundation Institute for Pediatric Research, Children’s Hospital of Pittsburgh of University of Pittsburgh Medical Center (UPMC), Pittsburgh, Pennsylvania, USA.; Department of Biostatistics, University of Pittsburgh, Pittsburgh, Pennsylvania; Acute and Tertiary Care Department, University of Pittsburgh School of Nursing, Pittsburgh, Pennsylvania.; Department of Medicine, University of Pittsburgh Medical Center, Pittsburgh, Pennsylvania

**Author notes:** Authors contributed equally to this work (Nolan Priedigkeit, Rebecca J. Watters). Shared senior authorship. Corresponding Author: Adrian V. Lee, Ph.D., Magee Women’s Research Institut, 204 Craft Avenue (Room A412), Pittsburgh, PA 15213, http://www.upci.upmc.edu/wcrc.

**Keywords:** Breast cancer, estrogen receptor, bone metastasis, RNA-seq, FFPE, decalcification, PAM50, tumor profiling, exome capture, cancer genomics, *RBBP8*

## Abstract

Bone metastases (BoM) are a significant cause of morbidity in patients with Estrogen-receptor (ER)-positive breast cancer, yet characterizations of human specimens are limited. In this study, exome-capture RNA-sequencing (ecRNA-seq) on aged (8-12 years), formalin-fixed paraffin-embedded (FFPE) and decalcified cancer specimens was first evaluated. Gene expression values and RNA-seq quality metrics from FFPE or decalcified tumor RNA showed minimal differences when compared to matched flash-frozen or non-decalcified tumors. ecRNA-seq was then applied on a longitudinal collection of 11 primary breast cancers and patient-matched *de novo* or recurrent BoM. BoMs harbored shifts to more Her2 and LumB PAM50 intrinsic subtypes, temporally influenced expression evolution, recurrently dysregulated prognostic gene sets and altered expression of clinically actionable genes, particularly in the CDK-Rb-E2F and FGFR-signaling pathways. Taken together, this study demonstrates the use of ecRNA-seq on decade-old and decalcified specimens and defines expression-based tumor evolution in long-term, estrogen-deprived metastases that may have immediate clinical implications.

**Grant Support:** Research funding for this project was provided in part by a Susan G. Komen Scholar award to AVL and to SO, the Breast Cancer Research Foundation (AVL and SO), the Fashion Footwear Association of New York, the Magee-Women’s Research Institute and Foundation, and through a Postdoctoral Fellowship awarded to RJW from the Department of Defense (BC123242). NP was supported by a training grant from the NIH/NIGMS (2T32GM008424-21) and an individual fellowship from the NIH/NCI (5F30CA203095).

**Conflicts of Interest Disclosure:** No relevant conflicts of interest disclosed for this study.

**Author Contributions:** Study concept and design (NP, RJW, SO, AVL); acquisition, analysis, or interpretation of data (all authors); drafting of the manuscript (NP, RJW, SO, AVL); critical revision of the manuscript for important intellectual content (all authors); administrative, technical, or material support (PCL, AB, RB, KRW, WH, JK, MR, ZF, AMB).

## INTRODUCTION

Bone metastases (BoM) occur in approximately 65-75% of breast cancer patients with relapsed disease, resulting in significant comorbidities such as fractures and chronic pain^1^. Following colonization to the bone, breast cancer cells exploit the local microenvironment by activating osteoclasts, which in turn provides proliferative fuel for tumor cells^2^. This process is targeted clinically using anti-osteoclast agents such as bisphosphonates and RANKL inhibitors, yet these therapies do not confer significant survival benefits^3^.

Importantly, the majority of breast cancers that metastasize to bone are estrogen receptor (ER)-positive and present clinically in the context of long-term endocrine therapies such as selective estrogen receptor modulators and aromatase inhibitors^4^. *In vivo* models of BoM have unfortunately been somewhat restricted to ER-negative disease due to the more indolent characteristics of ER-positive cell lines^5^. Molecular characterizations of ER-positive specimens that have recurred in an estrogen-deprived system, which represents the major burden of breast cancer BoM, are thus essential to reinforce the significant scientific contributions made using *in vivo* bone metastasis models^6–9^. Nonetheless, datasets are currently limited, in part due to the practical difficulties of obtaining and processing human BoM specimens^10^.

Large-scale molecular characterizations of patient-matched samples—primary tumors and synchronous or asynchronous matched metastases—show that metastatic lesions acquire features distinct from primary tumors that are either clinically actionable or confer therapy resistance^11–13^. Indeed, current treatment guidelines in breast cancer recommend a biopsy to guide therapy in advanced disease if possible^14^. Unfortunately, BoM often undergo harsh decalcification procedures with strong acids to eliminate calcium deposits prior to specimen sectioning. Decalcification degrades nucleic acids and can alter results of immunohistochemistry^15–17^. Furthermore, formalin-fixed paraffin embedding (FFPE)—often performed in concert with decalcification—causes severe degradation and hydrolysis of RNA^18^. In light of this, new capture-based methods of nucleic acid sequencing on aged FFPE specimens have shown efficacy in identifying DNA variants and even guiding care in academic centers^19–21^. Exome-capture RNA-sequencing (ecRNA-seq) is less well characterized in aged tumor samples, although recent studies on FFPE specimens have shown promising expression correlations with flash-frozen tissues^22–24^.

Because of the untapped potential of archived, decalcified BoM specimens, the burden of BoM in breast cancer patients and the lack of long-term endocrine treated tumor datasets, the performance of ecRNA-seq from decade-old, degraded and decalcified tumor samples was first assessed. Following this evaluation, ecRNA-seq was then applied to a collection of 11 ER-positive patient-matched primary breast cancers and bone metastases to define transcriptional evolution in breast cancer cells following metastatic colonization in the bone and years of endocrine therapy.

## RESULTS

### ecRNA-sequencing of aged and decalcified breast cancers

To determine the feasibility of sequencing an aged, FFPE and decalcified tumor cohort, ecRNA-seq on two separate sample sets was performed. The first sample set included four cases of primary breast tumors that at the time of resection, were split in two. One part was flash-frozen and stored at −80 C and the other tumor section was formalin-fixed paraffin embedded and stored at room temperature. Storage times ranged from 8.2 to 12.3 years. Post-alignment RNA-sequencing QC analyses showed differences in GC content and insert size, yet gene body coverage and transcript diversity assignments were largely similar (Figure 1A). After quantifying and normalizing gene abundances, expression correlations between frozen and FFPE matched samples were assessed using log2normCPM values. *Pearson r* correlations ranged from 0.929 to 0.963, with an average correlation of 0.953 (Figure 1B). The same analysis was performed using a second sample set of matched FFPE-decalcified and FFPE-non-decalcified samples. Again, no concerning deviations in RNA-seq quality metrics were observed between the two differently processed sample groups (Figure 1C) and Pearson *r* expression correlations ranged from 0.936 to 0.969 (Figure 1D). Furthermore, correlation matrices of the two sample sets showed matched tumor sample expression values were more similar to each other than expression values from tumors with equivalent processing and storage (Supplementary Figure 1). Full RNA-seq metrics from the QC analysis did reveal differences in some metrics between FFPE and flash-frozen tissue (i.e. splice junction loci number), that may be informative for other applications such as indel mutation calling or isoform detection (Supplementary Data S1 and S2). In summary, ecRNA-seq shows outstanding quality metrics for analysis of aged FFPE and decalcified bone metastases samples.

**Figure 1:**
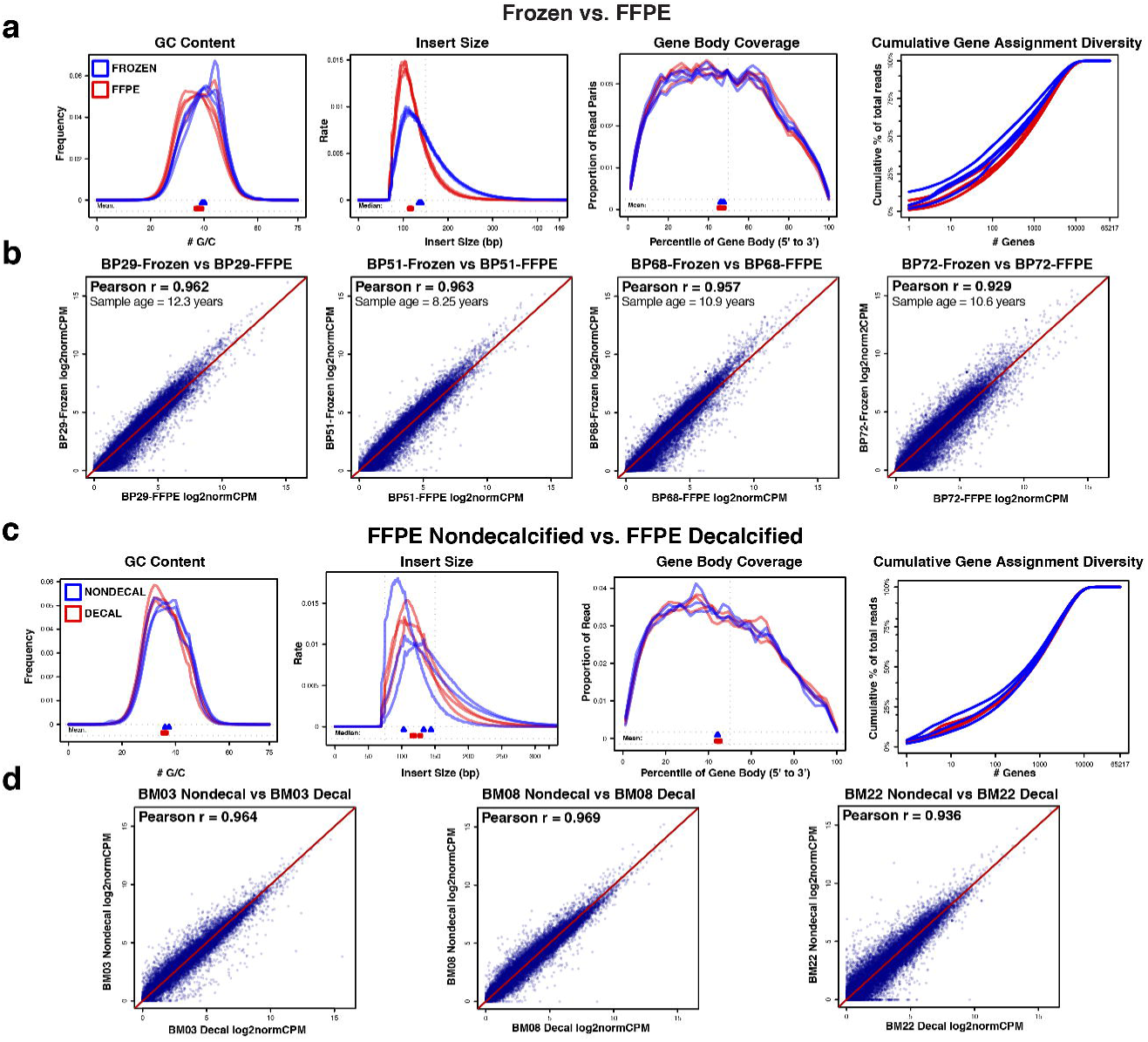
Exome-capture RNA-sequencing of aged, FFPE and decalcified tumors. **(A)** RNA-seq quality metrics (GC content, insert size, gene body coverage and cumulative gene assignment diversity) of aged and tumor-matched FFPE and flash-frozen (FF) sample; FF samples in blue, FFPE samples in red. **(B)** Expression value correlations between four sets of matched tumor samples (FF vs. FFPE) along with Pearson *r* correlations and sample ages. **(C)** RNA-seq quality metrics of matched nondecalcified and decalcified samples; non-decalcified samples in blue, decalcified samples in red. **(D)** Expression correlations between three sets of matched tumor samples (non-decalified vs. decalcified) along with Pearson *r* correlations.

### ecRNA-seq of breast cancer bone metastases

Following the validation of ecRNA-seq, a cohort of 11 ER-positive patient-matched primary tumors and BoMs was acquired through the University of Pittsburgh Health Science Tissue Bank (Table 1, Supplementary Data S3). Abstracted clinical records showed that nearly all patients (10/11) were documented as having received adjuvant endocrine therapy, and bone metastasis free survival ranged from 0 (de novo bone metastasis) to greater than 5 years with the most common site of bone metastasis being the vertebral column. ecRNA-seq was performed on the 22 samples yielding an average readcount of 58,294,593 and an average *Salmon* transcript mapping rate of 92.6% (Supplementary Data S4). Consistent with the initial quality control studies above, quality metrics on these samples showed consistent gene body coverage, GC content, insert sizes and transcript diversity regardless of decalcification status (Supplementary Figure 2, Supplementary Data S5). Furthermore, since samples within the cohort had been surgically excised and banked many years apart, all paired specimens underwent an analysis of shared variants, which confirmed tumor pairs were patient-matched (Supplementary Figure 3).

**Table 1:** Abridged clinicopathological features of patient-matched primary and bone metastasis tumor cohort^*¥*^.

| Case | Age at Dx | Histologic Subtype | Pathological Stage | ER Prim | PR Prim | HER2 Prim | BoM Location | BoM Decal | Endocrine Tx | HER2 Tx | Radio Tx | Chemo Tx | BMFS | OS |
| --- | --- | --- | --- | --- | --- | --- | --- | --- | --- | --- | --- | --- | --- | --- |
| **17** | 54 | IDC | IIIA | Pos | Pos | Neg | Ileum | Yes | Yes | No | Yes | Yes | 24 | 46 |
| **19** | 50 | IDC w/ lobular features | IV | Pos | Pos | Neg | Vertebra | No | Yes | No | Yes | No | 0 | 75 |
| **22** | 60 | IDC | IIA | Pos | Pos | Neg | Femur | No | Yes | No | Yes | Yes | 18 | 37 |
| **31** | 59 | IDC w/ lobular features | IIB | Pos | Pos | Neg | Vertebra | Yes | Yes | No | Yes | Yes | 43 | 55 |
| **34** | 38 | IDC | IIIA | Pos | Pos | Neg | Vertebra | Yes | Yes | No | Yes | Yes | 65 | 130 |
| **43** | 65 | IDC | IV | Pos | Pos | Neg | Vertebra | Yes | Yes | No | Yes | No | 0 | 54 |
| **44** | 56 | IDC | IA | Pos | Pos | Pos | Femur | No | NA | Yes | Yes | Yes | 23 | 42 |
| **48** | 49 | ILC | IIIC | Pos | Pos | Neg | Vertebra | No | Yes | No | Yes | Yes | 28 | 68 |
| **55** | 56 | IDC | IV | Pos | Pos | Neg | Femur | No | Yes | No | NA | No | 0 | 137 |
| **60** | 44 | IDC | IIB | Pos | Pos | Neg | Sacrum | Yes | Yes | No | Yes | Yes | 46 | 53 |
| **A25** | 39 | IDC | IIIA | Pos | Pos | Neg | Femur | Yes | Yes | Yes | Yes | Yes | 38 | 57 |

<sup>‡</sup>*Abbreviations:* Dx, diagnosis; Tx, therapy; ER, estrogen receptor; PR, progesterone receptor; HER2, human epidermal growth factor receptor 2; IDC, invasive ductal carcinoma; ILC, invasive lobular carcinoma; BoM, bone metastasis; BMFS, bone metastasis free survival; OS, overall survival

### Clustering and temporal expression shifts

Unsupervised hierarchical clustering of patient-matched pairs revealed that decalcification of BoMs did not produce independent clades, with 5 of 11 BoM clustering in the same doublet clade as their matched primary (denoted with * in Figure 2A). Notably, 3 of the 5 doublet clustering cases were de novo metastases. Discrete PAM50 intrinsic subtype assignments were identical in 6 of 11 pairs. 2 pairs switched from LumA to LumB in the metastasis, 1 pair from LumB to LumA, 1 pair from LumB to Her2 and another was classified as Normal subtype in the primary tumor and LumB in the BoM (Figure 2B). To obtain more granularity than discrete PAM50 calls, probability scores for each PAM50 subtype were assigned (Figure 2B and Supplementary Data S6). Her2 and LumB profile gains (defined as a probability gain of >10% in a matched BoM) were the most common— being observed in 4 of 11 cases (Figure 2B). Given observed shifts in expression profiles of bone metastases and doublet clustering of de novo bone metastases, temporal influence on transcriptional evolution was analyzed. Pearson *r* correlations between each patient-matched pair using log2normCPM expression values were utilized as a metric for transcriptional similarity. Expression pair similarity was significantly correlated (Pearson *r* = −0.864, *p-value* < 0.001) with time from primary tumor diagnosis to bone metastasis (Figure 2C).

**Figure 2:**
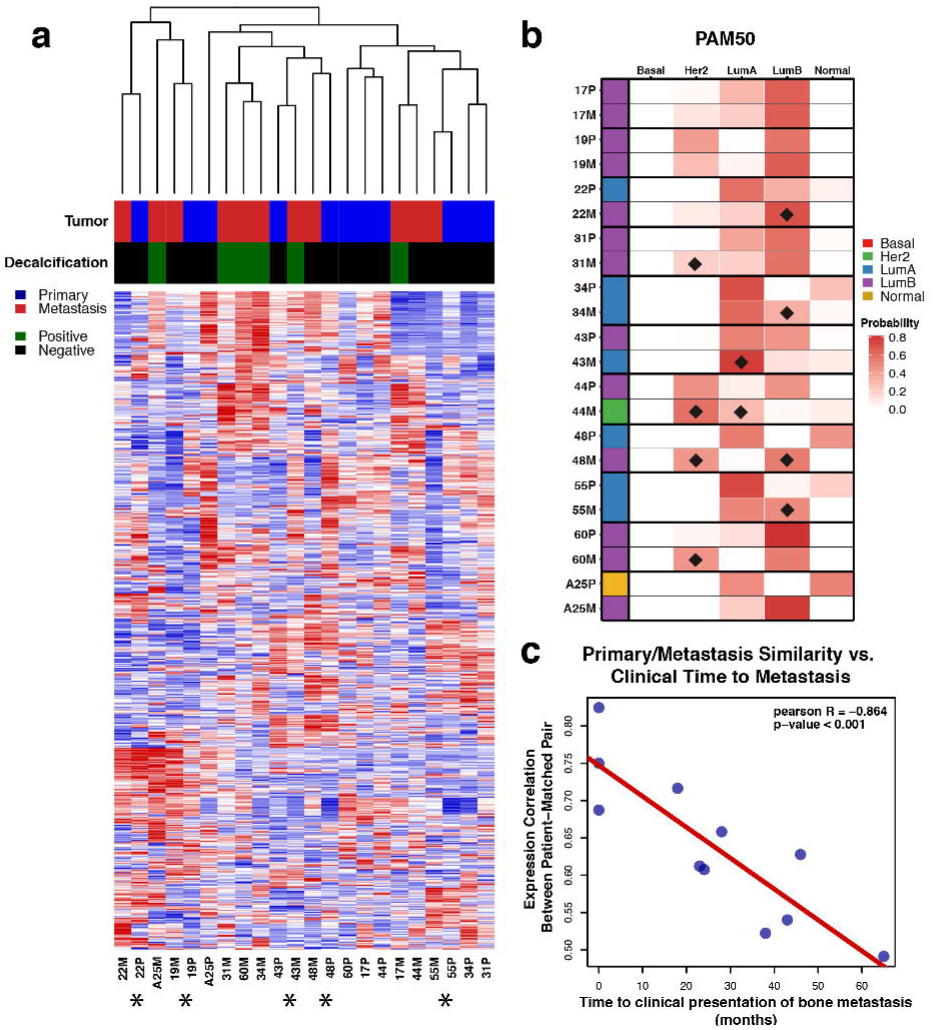
Unsupervised clustering, intrinsic subtype shifts and temporal evolution of ER-positive bone metastases. **(A)** Unsupervised hierarchical clustering heatmap (red = high relative expression, blue = low relative expression) of patient-matched pairs using the top 5% most variable genes (n = 1096) across the cohort. Tumor (primary in blue, metastasis in red) and decalcification status (positive in green, negative in black) indicated. Asterisks below heatmap designate patient-matched pairs that cluster in a single doublet clade. **(B)** Discrete PAM50 assignments (red = basal, green = HER2, blue = LumA, purple = LumB, yellow = Normal) and PAM50 probabilities for patient-matched pairs. PAM50 probability shifts in metastases (if greater than 10%) are marked with a black diamond. **(C)** Correlation of patient-matched tumor expression similarity versus clinical time to metastasis with Pearson *r* value and correlation p-value.

### Differentially expressed genes in bone metastases

To determine genes consistently up- or downregulated in bone metastases, a paired DESeq2 differential gene expression analysis was performed. 207 genes were differentially expressed (FDR adjusted *p-value* < 0.10)—80 genes with increased and 127 genes with decreased expression in bone metastases (Figure 3A, Supplementary Data S7). Gene ontology analysis was performed to determine biological processes represented in the up- and downregulated gene sets. Generally, genes within osteogenic programs showed the most significant increases in expression while muscle-related, adhesion and motility gene sets were found to be significantly lost in bone metastases (Figure 3A, Supplementary Data S8, Supplementary Figure 4). Given that a subset of these genes may be mediating therapy resistance and/or distant metastases, single sample gene set enrichment analysis (ssGSEA) scores^25^ were calculated using tumor expression data from patients with long-term outcomes in METABRIC^26^. Two separate gene lists were created to build the signatures—representing the most significantly upregulated (boneMetSigUp) and downregulated (boneMetSigDown) genes in bone metastases (Supplementary Data S9). Tumors intrinsically expressing higher boneMetSigUp and lower boneMetSigDown ssGSEA scores conferred worse (log-rank *p-value* < 0.001) disease-specific survival outcomes (Figure 3B). To increase the power of discerning gene expression effects due to long-term estrogen deprivation, a differential gene expression analysis was performed excluding the treatment-naïve, de novo bone metastases. This yielded a list of 612 differentially expressed genes (Supplementary Data S10), some of which were not detected as differentially expressed with treatment-naïve de novo bone metastasis cases included.

**Figure 3:**
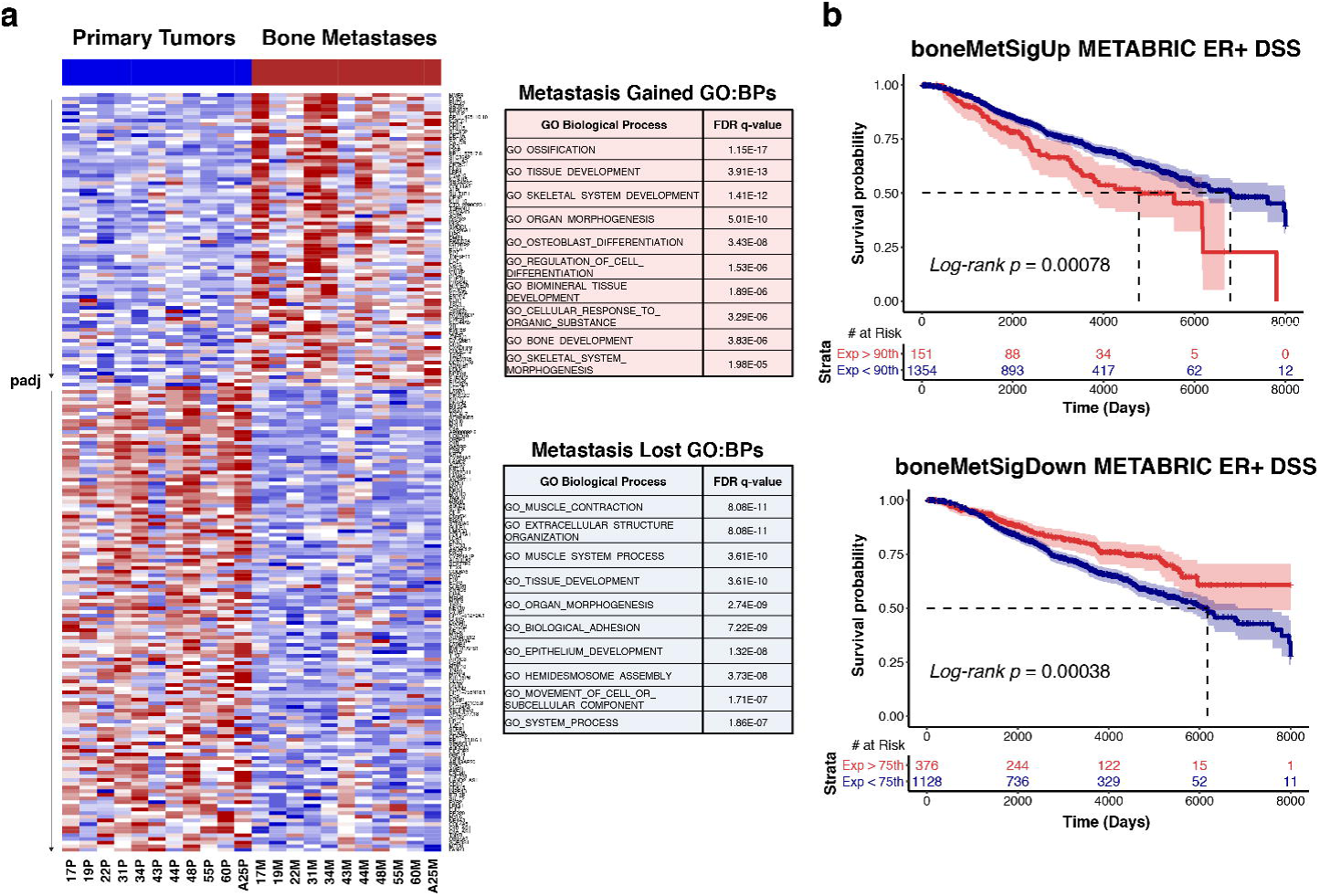
Differentially expressed genes in patient-matched bone metastases. **(A)** Left, heatmap (red = high relative expression, blue = low relative expression) of log2normCPM values from 207 differentially expressed genes (FDR adjusted p-value < 0.10) between primary tumors and patient-matched bone metastases. Heatmap is segregated into two sections; genes with log2FoldChange > 0 on top and genes with log2FoldChange < 0 on bottom. Each section is gene-sorted by adjusted p-values. Right, Gene Ontology: Biological Process gene overlap analysis for genes with significant expression gains (top, red) and losses (bottom, blue) in bone metastases. Top 10 pathways are shown alongside FDR adjusted q-values. **(B)** Disease-specific survival outcome differences in ER-positive METABRIC tumors using boneMetSigUp (top) and boneMetSigDown (bottom) expression scores as strata. 95% confidence intervals are highlighted along with log-rank p-values and associated risk tables.

### Dysregulated gene sets and RBBP8 expression loss

To determine pathway level changes in breast cancer bone metastases, a pre-ranked GSEA was performed. All genes were ranked by DESeq2 calculated log2 fold-changes (metastasis vs. primary, Supplementary Data S11) and then analyzed for enrichments using Molecular Signature Database (MsigDB) gene sets (http://software.broadinstitute.org/gsea/msigdb, H:Hallmark gene sets, C6: Oncogenic signatures)^27^.This yielded several significantly metastasis-enriched and metastasis-diminished gene sets (FDR *q-val* < 0.10, Supplementary Data S12). The three most significantly enriched gene sets in metastases involved E2F transcription factor targets, genes mediating the G2M checkpoint and an experimental perturbation gene set consisting of genes up-regulated with knockdown of *RBBP8* in a breast cell line (Figure 4A). Other upregulated gene sets included hedgehog signaling and gene sets associated with Rb loss and KRAS gains. The three most significantly negatively correlated gene sets consisted of an NFKb/TNF gene set, genes involved in epithelial mesenchymal transition (EMT) and an embryonic development gene set. We further interrogated *RBBP8* due to it being the most significant gene set enriched in bone metastasis. As predicted by the enrichment, bone metastases carried significant *RBBP8* expression loss (Wilcoxon-signed rank *p-value* = 0.02), with 5 of 11 metastases [45%] having at least a 2-fold decrease in expression versus patient-matched primaries (Figure 4B). Tumors intrinsically expressing lower levels of *RBBP8* showed worse disease-specific and bone metastasis-free survival outcomes (Figure 4C).

**Figure 4:**
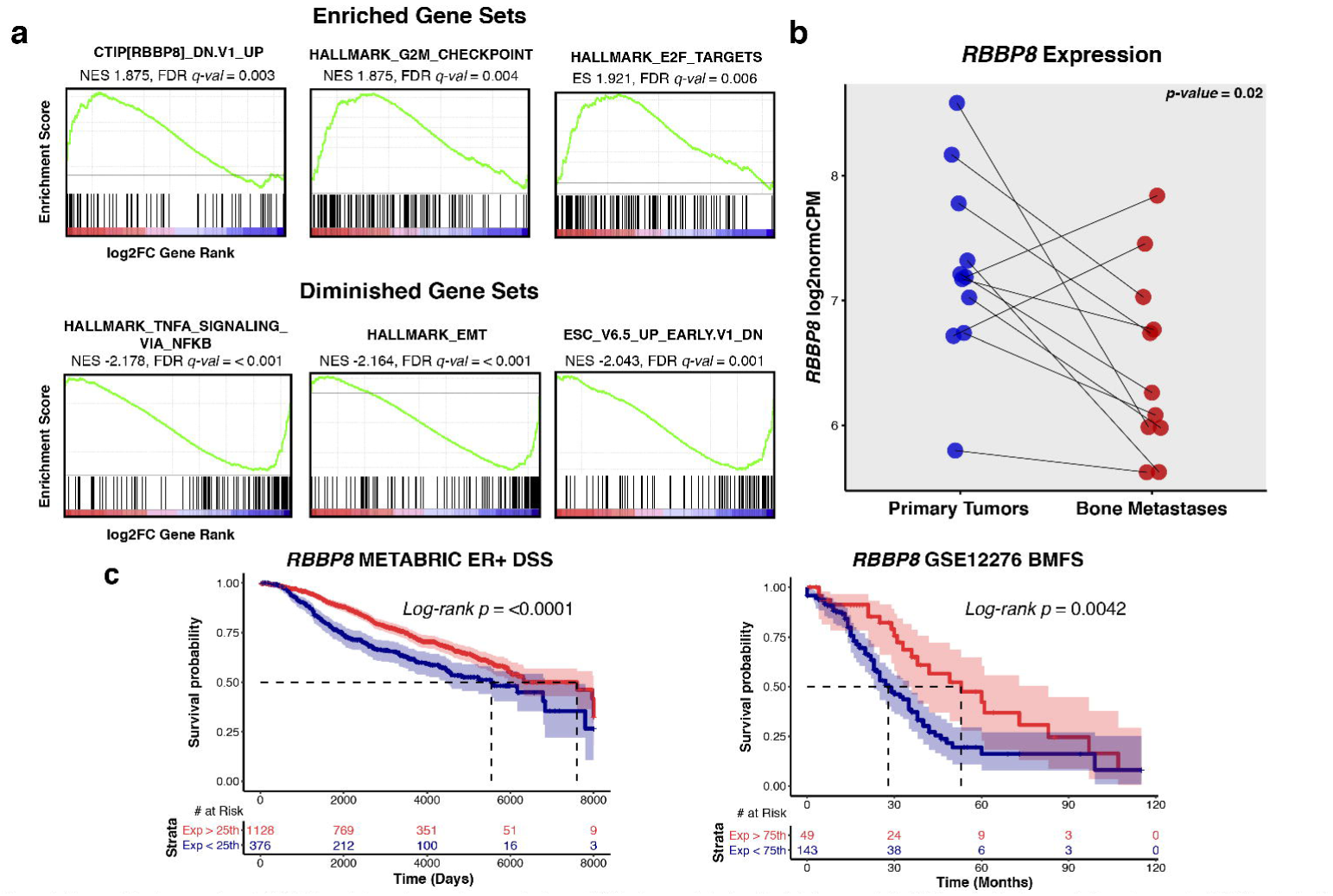
Dysregulated gene sets and RBBP8 loss in breast cancer bone metastases. **(A)** Top three enriched and depleted gene sets (by FDR q-value) in bone metastases from ranked GSEA analysis. Gene list ranking was performed using log2FoldChange values from DESeq2 differential expression output, where a positive log2FoldChange represents increased expression in metastasis (red) and a negative log2FoldChange represents decreased expression in metastasis (blue). Green line shows running enrichment score as algorithm walks down the ranked gene list. Black vertical lines below curve show where genes within the query gene set are represented in the ranked list. Normalized enrichment score (NES) and FDR q-values are noted below gene set names. **(B)** *RBBP8* expression values (log2normCPMs) in primary tumors (blue) and bone metastasis (red). Pairs are connected with a line and Wilcoxon signed-rank *p-value* is shown. **(C)** Disease-specific survival outcome differences in ER-positive tumors (METABRIC) and bone metastasis free survival differences (GSE12276) using normalized *RBBP8* expression values as strata. 95% confidence intervals are highlighted along with log-rank p-values and risk tables.

### Expression gains and losses in clinically actionable genes

Because of the observed acquisition of clinically actionable targets reported in other studies of paired primary and recurrent tumors^12,13^, a paired expression analysis to define clinically actionable expression changes in ER-positive bone metastases was performed (Supplementary Data 13). Using stringent, case-informed cutoffs for expression alterations (Supplementary Figure 5), the most common expression losses in bone metastases were *PIK3C2G* [8 of 11, 73%], *ESR1 [7 of 11, 64%] and TUBB3 [6 of 11, 55%]* (Figure 5A and Supplementary Figure 6). Other notable losses included *GREM1, PTPRT, CDKN2A, KIT* and *GATA3*. The most recurrent expression gains were *FGFR3* [7 of 11, 64%], *EPHA3 and PTPRD* [6 of 11, 55%]. *PDGFRA, PTCH1, ALK, HGF, FGFR1* and *FGFR4* also showed highly recurrent gains (Figure 5B). Interestingly, some expression gains were absent in *de novo* bone metastasis cases (Cases 19, 53 and 55) yet highly recurrent in long-term endocrine-deprived cases (*EPHA3, PTPRD, PDGFRA, PTCH1*), suggesting clinically actionable, treatment-driven gains in endocrine-resistant breast cancer recurrences.

**Figure 5:**
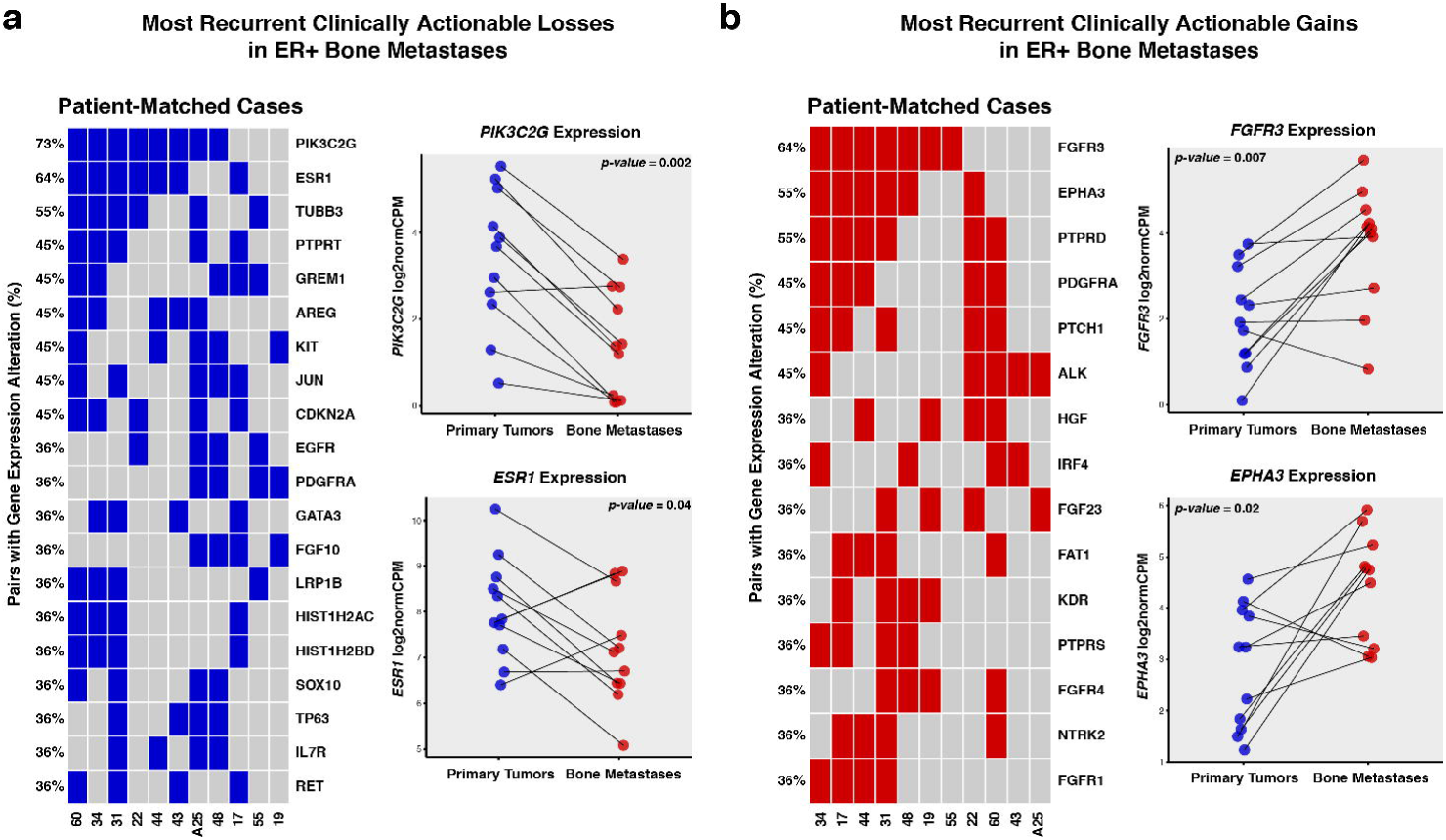
Recurrent, clinically actionable expression gains and losses in ER-positive bone metastasis. **(A)** Recurrent expression alteration losses, ranked by frequency, for each patient-matched case (columns). Each blue tile represents a bone metastasis with a lower log2FoldChange vs. its matched primary than the case-specific expression loss threshold. Expression values (log2normCPMs) for most recurrent losses (*PIK3C2G, ESR1*) are pair plotted with corresponding Wilcoxon signed-rank test *p-values* noted. **(B)** Recurrent expression alteration gains, ranked by frequency. Red tiles represent bone metastases with higher log2FoldChange than the case-specific expression gain thresholds. The two most recurrent expression gains (*FGFR3, EPHA3*) are also plotted.

## DISCUSSION

Bone is the most common site of distant recurrence for patients with ER-positive breast cancer, yet comprehensive sequencing datasets of endocrine therapy treated, metastatic samples are currently limited. This is in part due the challenge of obtaining tissue, and degradation of nucleic acids caused by decalcification. In this study, we found that aged FFPE and FFPE-decalcified tumors showed highly similar transcript quantification values as matched flash-frozen and FFPE-non-decalcified tumors. As a proof-of-concept, we then applied ecRNA-seq to a cohort of patient-matched primary and bone metastases collected over a period of five years. We identified subtle shifts in intrinsic subtypes and found a strong temporal influence on transcriptional evolution in breast cancer recurrences. Furthermore, we created several differentially expressed gene sets/signatures that are prognostic and point towards acquired *RBBP8* loss, CDK-Rb-E2F and FGFR pathway gains as mediators of ER-positive breast cancer progression. Lastly, we found bone metastases commonly gain or lose expression in clinically actionable genes, which may be distinct from primary tumors.

ecRNA-seq is an effective method for quantifying expression on aged, FFPE and decalcified tumor specimens. Previous work has assessed nucleic acid amplification success, DNA-sequencing and RNA integrity metrics using decalcified samples^17,28,29^; however, a comprehensive analysis of RNA-sequencing, to our knowledge, has not yet been performed. Consistent with only very minor differences between GC content, insert sizes and other QC metrics, gene expression values between aged matched FFPE/flash-frozen and FFPE-decalcified/FFPE-non-decalcified tumors are highly correlated (Pearson *r* range 0.929 – 0.969). This study reinforces and should encourage the use of capture-hybridization approaches to sequence RNA from retrospectively collected, low yield, highly degraded and decalcified archival specimens (Supplementary Data S15)^22–24^. Expanding sample sets and modalities for genome-wide characterization, especially for rare specimen cohorts that may be impractical to obtain prospectively in large numbers, will accelerate translational discoveries.

Given promising results from our evaluation, we applied ecRNA-seq in a proof-of-concept effort to characterize the transcriptome of 11 archival patient-matched ER-positive primary and recurrent metastases— 3 cases having treatment-naïve, *de novo* bone metastases and 8 recurrent cases harboring long-term endocrine-therapy treated metastases. In the recurrent cases, bone metastasis-free survival ranged from 18 to 65 months. Despite a large portion of the bone metastases being decalcified, global transcriptome QC metrics showed similar features (i.e. GC content, insert sizes, gene body coverage and transcript assignment diversity) and no outliers. Consistent with this, unsupervised hierarchical clustering showed no distinct clusters of decalcified samples, with 5 bone metastases clustering in the same doublet clade as their patient-matched primary breast cancer. Interestingly, 3 of these doublet clustering pairs were clinically *de novo*, treatment naïve bone metastases, implying limited transcriptional evolution from the primary tumor in synchronous metastases. This was further corroborated with a striking negative correlation between patient-matched expression similarity and time to bone metastasis, suggesting metachronous metastases that clinically present later in their treatment course are more dissimilar from their derived primary lesions. Intrinsic subtyping revealed 5 of the 11 cases changed PAM50 subtypes, with 3 cases switching to LumB in the metastasis and another switching to Her2. Subtle Her2 and LumB profile shifts were also the most common when observing continuous PAM50 probability scores, even in samples that remained concordant in their discrete PAM50 assignments. A recent, targeted expression study analyzed PAM50 assignments in 123 matched breast cancer metastases and the authors found similar frequencies of LumB and Her2 acquisitions in ER-positive metastatic tumors^30^. Given this transcriptional evolution to more LumB and Her2 profiles, a thoughtful reevaluation of therapy selection in the advanced and perhaps the adjuvant setting may be necessary—especially considering HER2-targeted therapies are generally reserved for patients with HER2-positive primary disease.

We found 207 genes to be differentially expressed between primary tumors and patient-matched bone metastases. The top upregulated genes belonged to osteogenic gene sets—*BGLAP, RANKL, PTH1R* all showing significant expression gains—and supports *in vivo* modelling observations of breast cancer osteomimicry and hijacking of the bone microenvironment^31^. Downregulated gene sets included genes involved in broad categories such as cellular adhesion, hemidesmosome assembly and epithelium development, pointing towards specific biological programs lost following metastatic colonization. Moreover, when either the upregulated or downregulated genes are expressed coordinately in primary tumors, we found that they confer worse and better outcomes respectively in ER-positive tumors, suggesting some tumors may develop these transcriptional programs early in their evolution. Lastly, a differential expression analysis between endocrine naïve primary tumors and long-term endocrine treated bone metastases identified a larger list of differentially expressed genes. Importantly, known mediators of endocrine resistance are represented in the list, including dysregulated expression of *Wnt* family members^32^, expression gains in FGFR1^33^, *FOXC1^34^* and loss of *ESR1* expression^35^. Notably, many of these genes do not overlap with the differential expression analysis that included the de novo metastases, suggesting expression alterations specific to late recurrent therapy-treated tumors. This non-overlapping gene set included a greater than 2-fold average expression gain of *ABCG2* in therapy-exposed metastases—a multidrug resistance protein shown to be active in breast cancer^36,37^—and loss of *CDKN2A*. CDKN2A encodes *p16*, a negative regulator of CDK4/CDK6 and is located on a common somatically deleted region (9p21) in cancer^38^. Given recent success of CDK4/CDK6-inhibiting compounds (palbociclib and ribociclib) in treating ER-positive breast cancers, this recurrent, acquired, metastatic-specific loss of *CDK2NA* is a clinically important observation^39–41^.

Following significant gene-level changes, a gene set enrichment analysis defined enriched and diminished pathways in breast cancer bone metastases. Enriched genes included those involved in G2M checkpoint and E2F targets. Consistent with the observed LumB enrichments, breast cancer cells appear to develop a more proliferative phenotype following bone colonization and the strong enrichment of E2F signature in metastatic disease again highlights the CDK-Rb-E2F pathway as a potential actionable target. Interestingly, another study that utilized a targeted gene expression platform found proliferative gene signatures in ER-positive metastases may be more accurate at predicting overall survival than signatures in the primary tumor^30^. A survival analysis for this work was impractical given the small set of patient-matched pairs, but future metaanalyses are warranted to determine if gene expression signatures in metastases are better predictors of overall survival in the advanced setting, especially given the significant transcriptomic shifts observed in this study.

The most significant gene set enriched in bone metastasis was an experimental perturbation gene set involving the knockdown of the tumor suppressor RBBP8^42^. *RBBP8* (also known as CtIP) binds directly to Rb, mediates cell cycle regulation, helps maintain genomic stability and loss of *RBBP8* incurs tamoxifen resistance and sensitizes breast cancer cells to PARP inhibition *in* vitro^43–46^. Concordant with the GSEA analysis, bone metastases have significant expression loss of *RBBP8*, with 45% of cases showing a greater than 2-fold decrease in expression. We found low *RBBP8* expression in ER-positive tumors confers poorer disease-specific survival and bone metastasis-free survival outcomes. These observations point to *RBBP8* loss in metastatic breast cancers as being a prime, perhaps therapeutically relevant candidate for further preclinical investigations.

Lastly, considering we have previously shown that brain metastases acquire highly recurrent gains in clinically actionable genes^13^, particularly in HER2, we analyzed the same set of genes in bone metastases. All tumors harbored significant gains and losses, some of which were highly recurrent. *PIK3C2G*, a relatively uncharacterized gene in the PI3K pathway, was the most recurrent gene expression loss. Other notable losses included *ESR1, CDKN2A* and *GATA3*—genes that have already been implicated in endocrine therapy resistance in experimental models. Intriguingly, *GATA3* is one of the most recurrently mutated genes in breast cancer, being particularly enriched in ER-positive disease^47^. Moreover, *GATA3* inhibits breast cancer metastasis in various model systems and given losses of *GATA3* in ER-positive bone metastases are common, further evaluation of *GATA3* as a potentially targetable breast cancer metastasis suppressor gene should be encouraged^34,48,49^. Metastatic gains included FGFR family members (*FGFR3, FGFR4, FGFR1*), *ALK* and *KDR*—all protein products having small molecules currently in clinical trials. Interestingly, some highly recurrent expression gains (i.e. *EPHA3, PTPRD, PDGFRA, PTCH1*) were exclusive to long-term endocrine treated bone metastases suggesting them as prime, clinically actionable candidate mediators of therapy resistance. Collectively, these observations provide yet further evidence of acquired transcriptional programs in metastatic lesions and suggests that precision care in breast cancer should be informed by molecular features of advanced tumors in order to not miss metastatic dependencies acquired in advanced disease.

Although this study points towards ecRNA-seq as being a viable option to characterize the transcriptome of archived, decalcified specimens, there are limitations. Firstly, multiple methods are used for decalcification with varying effects on nucleic acids and we were unaware of this information for the profiled specimens, as it is rarely recorded in clinical notes^17^. Secondly, in primary versus metastatic expression studies, it is difficult to deconvolute expression contributions from tumors versus the altered microenvironment of the distant organ site. To limit these artifacts in this study, regions of high tumor cellularity in the bone metastasis were cored by a trained molecular pathologist for RNA extraction, which is corroborated by RNA-seq derived tumor purity estimates—as no significant tumor purity differences between primary and metastatic tumors (Supplementary Data S15) were observed^50^. Nonetheless, single-cell sequencing approaches of metastatic tumors will be essential to bring cell-level resolution to transcriptional studies of metastatic tumors. Novel computational methods that deconvolute heterogeneous sample sets, until single-cell sequencing becomes more widely adopted, will also be essential^51–53^. All of this withstanding, features of the data are encouraging such as patient-matched tumors clustering together, intuitive PAM50 assignments, corroboration of other groups’ findings and treatment-specific gains and losses. Finally, a limitation of this study is the small sample size. Hopefully, these results will encourage the use of ecRNA-seq to transcriptionally profile other highly degraded samples and begin a collection of genomic data from metastatic or rare tissues for integration. Importantly, de-identified clinical data should be provided alongside the sequencing, as in this study, to allow more fluid merging of datasets and inspire clinical phenotype-driven analyses.

Taken together, this study both validates the use of ecRNA-seq to transcriptionally profile highly degraded RNA from decade-old and decalcified tumor specimens and defines multiple acquired and lost transcriptional programs in ER-positive bone metastases. We highlight acquired changes in the CDK-Rb-E2F and FGFR pathways, particularly relevant given the recent clinical use of CDK4/6 inhibitors, and point towards *RBBP8* as a particularly compelling candidate in breast cancer progression. We also find significant gains in clinically actionable genes that may have not been appreciated in primary tumors, reinforcing the need for longitudinal characterizations of cancer specimens to guide clinical care.

## METHODS

### Sample acquisition

Eleven sets of formalin-fixed paraffin-embedded (FFPE) primary breast tumors and patient-matched bone metastases (total of 22 samples) were obtained from the Health Sciences Tissue Bank, a certified honest broker facility at the University of Pittsburgh that maintains an IRB-approved protocol for collecting excess tissue and biological materials. A molecular pathologist reviewed hematoxylin and eosin slides from each sample and then subsequently cut 0.6-1 mm cores from the paraffin block exclusively from regions of high tumor cell purity for RNA extraction. De-identified clinical and biological data were collected under the approval of the University of Pittsburgh Institutional Review Board (Protocol numbers: PRO14040193 and PRO10050461).

### Tissue processing and RNA extraction

Tissues were digested over-night with shaking at 300 rpm at 56 °C in PKD buffer with the addition of proteinase K (Qiagen). RNA extraction was then performed with Qiagen’s FFPE RNeasy kit (Qiagen, Cat#73504) according to the manufacturer’s instructions under sterile RNase/DNase free conditions. RNA concentration was determined with the Qubit 3.0 Fluorometer (ThermoFisher Scientific). Quality RNA integrity number (RIN) scores and fragment sizes (DV200 metics) were obtained utilizing either the Agilent 2100 Bioanalyzer or the Agilent 4200 TapeStation.

### Exome-capture RNA-sequencing

Sequencing library preparation was performed using a minimum of 25 ng of RNA according to Illumina’s TruSeq RNA Access Library Preparation protocol. Indexed, pooled libraries were then sequenced on the Illumina NextSeq 500 platform with a High Output flow cell producing stranded, paired-end reads (2 X 75 bp). A target count of 50 million reads per sample was used to plan indexing and sequencing runs.

### RNA-sequencing expression quantification and normalization

RNA transcripts from paired-end FASTQ files were mapped and quantified using k-mer based lightweight-alignment with seqBias and gcBias corrections (Salmon v0.7.2, quasimapping mode, 31-kmer index built from GRCh38 Ensembl v82 transcript annotations)^54^. Transcript-level abundance estimates were collapsed to gene-level estimates using tximport2^55^. To filter out non- or low expressed genes, only genes harboring a TPM value of more than 0.5 in at least 10% of samples were considered. Gene-level counts or log2 transformed TMM-normalized CPM (log2normCPM) values were implemented for subsequent analyses.^56,57^.

### Expression correlations and RNA-seq quality assessment

Exome-capture RNA-seq was performed on two cohorts: 1) a set of four aged (ranging from 8 – 12 years) primary breast cancer specimens that at the time of surgical resection were split in half and either immediately embedded in optimal cutting temperature (OCT) compound and flash-frozen for storage at −80C, or formalin-fixed paraffin embedded (FFPE) and stored at room temperature. A second cohort consisted of three breast cancer bone metastases that at the time of resection were split in half and either decalcified or nondecalcified and processed to FFPE. These datasets were quantified and normalized as described above. Pearson *r* correlations between all samples were determined using log2normCPM values. Reads and mapping rates were obtained from *Salmon*. More detailed RNA-seq metrics were calculated and plotted using QoRTs (v1.1.8) following two-pass read alignment with STAR (v2.4.2a) for the 11 patient-matched cases^58,59^.

### tumorMatch patient-matched sample identifier

To confirm samples were patient-matched, variants from RNA-seq were called using *GATK’s Best Practices for variant calling on RNA-seq*^60^. Output .vcf files were then provided to *tumorMatch*, a custom *R* script that analyzes a pool of .vcf files and calculates the proportion of shared variants (POSV) between each .vcf. These proportion values were visualized using *corrplot* in *R*^61^.

### Unsupervised hierarchical clustering and intrinsic subtyping

Hierarchical clustering was performed using the heatmap.3 function (https://raw.githubusercontent.com/obigriffith/biostar-tutorials/master/Heatmaps/heatmap.3.R) in R on log2normCPM values of the top 5% most variable genes (defined by IQR) with 1 minus Pearson correlations as distance measurements and the “average” agglomeration method. PAM50 calls were generated using the *molecular.subtyping* function in *genefu* ^62^. A separate cohort of exome-capture RNA-sequencing expression data from primary tumors (n = 12 ER-negative, 9 ER-positive) was merged with the bone metastasis cohort to help account for test-set bias and increase the stability of the PAM50 assignments^63^. To call PAM50 subtypes, for each query sample in the bone metastasis cohort a random subset of primary tumor expression data was added to enforce a balanced distribution of ER-positive and ER-negative tumors. This was repeated 20 times and the discrete PAM50 subtype was designated as the mode of this 20-fold PAM50 assignment test while the final probability score was an average of all 20 probability scores from *genefu*.

### Differential gene expression

Salmon gene-level counts with effective lengths of target transcripts were used to call differentially expressed genes (DEGs) between primary tumors and bone metastases using DESeq ^64^. Given samples were patient-matched, a multi-factor design was implemented (∼Patient + Tumor [i.e. primary vs. metastasis]). Genes with an FDR adjusted p-value of less than 0.10 were assigned as differentially expressed. An unclustered heatmap using log2normCPM values from the 207 DEGs, first segregated by metastatic log2FoldChange gains and losses and then sorted by DESeq2 adjusted p-values, was created in R using heatmap.3. Differentially expressed genes within the *MsigDB* database that were gained or lost in bone metastases were separately interrogated for gene ontology (GO: Biological Process) enrichment by computing significant (top 10 gene sets) gene overlaps using the MsigDB online tool^27^.

### ssGSEA signatures and METABRIC survival analyses

Microarray expression along with disease-specific survival (DSS) data was obtained from the Molecular Taxonomy of Breast Cancer International Consortium (METABRIC) through Synapse (https://www.synapse.org/, Synapse ID: syn1688369), following IRB approval for data access from the University of Pittsburgh^26^. Normalized expression values from IHC-confirmed ER-positive tumors were used to develop a single-sample gene-setenrichment score (ssGSEA) for strongly DEGs (adjusted p-value < 0.05) between primary tumors and bone metastases^25^. 48 genes that carried positive log2FoldChange values and had a corresponding gene expression value in METABRIC were assigned to the “boneMetSigUp” signature; 74 genes with negative log2FoldChange values were assigned to the “boneMetSigDown” signature. A ssGSEA score for each sample from both gene sets was calculated using the ssGSEA method implemented in the *GSVA R* package^65^. Binary dichotomization of samples (low vs. high) based on ssGSEA signature score strata (10th, 25th, 50th, 75th, 90th percentiles) and log-rank testing were used to assess significant differences in DSS^66^. The strata with the most significant log-rank p-values were plotted using *survminer* from CRAN^67^.

### Ranked Gene Set Enrichment Analysis (GSEA)

To determine pathways significantly enriched or lost in breast cancer bone metastases versus patient-matched primaries, GSEA analyses were performed using gene sets with coordinately expressed genes representing specific biological and cancer-related pathways (MSigDB: H and C6 sets). Input into GSEA was a ranked list (DESeq2 log2FoldChange values) of 21,702 genes. Enrichment scores, significance values and plots were generated using default settings of the Broad Institute’s javaGSEA Desktop Application (v2.2.3).

### RBBP8 survival analysis

*RBBP8* expression was further interrogated and plotted using log2normCPM values from patient-matched. *RBBP8* expression influence on DSS in METABRIC ER-posiitve patients was interrogated as described above. RBBP8 expression influence on bone-met free survival (BMFS) was assessed by querying a GCRMA-normalized microarray expression dataset (GSE12276) from 204 primary tumors and associated survival data as described above^68^.

### Gains and losses in clinically actionable genes

Clinically actionable gene set was obtained using the Drug Gene Interaction Database (DGBIdB 2.0)^69^. Considering metastatic fold-change distributions calculated from log2normCPM values for all genes were slightly different for each case, stringent case-specific fold-change thresholds were used to transform continuous fold-change values into discrete “expression alterations.” More specifically, if the fold-change value for a clinically actionable *GENE_X* was greater than the 95^th^ percentile of all gene fold-change values in that case, *GENE_X* would be designated as a significant, case-specific expression gain. If the fold-change value for *GENE_Y* was lower than the 5^th^ percentile, *GENE_Y* was designated as a significant, case-specific expression loss (Supplementary Figure 6, Supplementary Data S13). After assigning discrete expression alteration calls to clinically actionable genes, data was visualized using the *oncoprint* function in *ComplexHeatmap*^70^.

### Statistical considerations

To determine differentially expressed genes between patient-matched primary tumors and bone metastases, *DESeq2* was used. *DESeq2* is designed for RNA-seq gene-based count abundance estimates and assigns differential expression *p-values* based on a negative binomial distribution. For Kaplan-Meier curves, the logrank test was used to determine statistically significant differences in event probabilities (i.e. death or time to metastasis) based on binary expression or signature strata. For single gene queries, paired Wilcoxon-signed ranked tests on log2normCPM values were used.

## Data availability

Comprehensive expression values for all samples will be deposited in the Gene Expression Omnibus (GEO). Raw sequencing data will be available upon request from authors and delegated in accordance to Institutional Review Board policies.

## Code availability

A collated version of code used to produce the major figures in this manuscript will be made publically available as performed for previous publications (https://github.com/npriedig/).

## SUPPLEMENTARY FIGURES AND LEGENDS ARE PROVIDED IN SEPARATE FILE

## Acknowledgements

This project used the University of Pittsburgh HSCRF Genomics Research Core and Health Sciences Tissue Bank, and the UPCI Tissue and Research Pathology Services supported in part by award P30CA047904. The authors would like to thank the patients who contributed samples to this study and Lori Miller (University of Pittsburgh), Alma E. Heyl (UPMC) and Jorge A. Rios (UPMC) for their efforts in collecting tissues.

